# Host plant nutrition drives fitness outcomes in the cactus specialist *Drosophila mettleri*

**DOI:** 10.1101/2025.09.09.675243

**Authors:** Lidane Noronha, Brian P. Lazzaro, Patrick M. O’Grady

**Affiliations:** Department of Entomology, Cornell University, Ithaca, New York, United States of America

**Keywords:** Ecological adaptation, Fitness consequences, Host plant specialization, Microbial interactions, Nutritional ecology, Plant-insect interactions

## Abstract

Organisms must navigate complex interactions with host plants, microbial communities, and environmental cues to ensure their survival and reproductive success when adapting to novel environments. Host plant specialists can be used to study how these interactions affect fitness due to their ecological constraints. In specialist species, such as cactophilic *Drosophila*, it remains unclear how feeding preferences, nutritional value of the substrate, and microbial interactions collectively shape fitness outcomes. We used diverse laboratory media formulations to examine the fitness implications in *Drosophila mettleri*, a species that exclusively uses Saguaro cactus (*Carnegiea gigantea*) as its natural host plant. We found that *D. mettleri* did not preferentially feed on one medium compared to others, nor on *C. gigantea* supplemented media compared to controls. Microbial communities, especially on banana media with *C. gigantea* additives, appear to improve developmental success and can influence *D. mettleri* post-pupation success. Taken together, our results highlight the behavioral and environmental interplay between host plant substrate and microbial communities.

## Introduction

Adapting to a novel host plant may require several steps, including tolerance to plant defensive compounds, developing a preference for the volatile and nutritional profile of a given host substrate, and subsequently becoming dependent on that host for metabolism and development (Zhang et al., 2015; Sharma et al., 2020). Host shifts can affect oviposition behaviour, morphology, interspecific competition, and, at times, can lead to specialization (Kibota & Courtney, 1991; Winkler & Mitter, 2008; Bouzas et al., 2021). Tripartite symbiosis between insects, plants, and microbes such as the cactus–*Drosophila*–yeast model system, has been used to examine the broad effects of host plant specialization (Fellows & Heed, 1972; Starmer et al., 1976; Starmer et al., 1982; Markow & O’Grady, 2008; Castrezana & Bono, 2012; Bouzas et al., 2021). However, little is known about how cactus and yeast act together to affect the fitness of *Drosophila* species.

Deserts exist on the edge of environmental extremes, making them a natural laboratory to study host shifts under climatic pressure (Franklin et al., 2016). The added layer of environmental stresses, such as scarcity of water and food sources, can lead to close evolutionary relationships between insects and their host plants, with microbes mediating these interactions (Lachance & Starmer, 2008; Lachance et al., 2013; Franklin et al., 2016). The Saguaro cactus, *Carnegiea gigantea*, is emblematic of the Sonoran Desert (*Saguaro - Saguaro National Park* (U.S. National Park Service, 2024). Saguaro produces alkaloid toxins that are toxic to most animals, except the few that have evolved mechanisms to detoxify the exudate (Fogleman, Heed, et al., 1982; Fogleman & Danielson, 2001). Two species of *Drosophila* endemic to the Sonoran Desert, *D. nigrospiracula* and *D. mettleri*, utilize necrotic Saguaro as an oviposition substrate (Castrezana & Bono, 2012), but feed on the yeast florae found in this environment (Fogleman, Starmer, et al., 1982). *Drosophila mettleri* oviposits in soil that has been soaked in necrotic cactus tissue (Heed 1977). High evaporation rates within the arid Sonoran Desert concentrate toxins in the soil as the exudate accumulates (Meyer & Fogleman, 1987), rendering this environment free of competition from other arthropods, including the closely related *D. nigrospiracula,* which oviposits in the necrotic Saguaro cactus tissue (Castrezana & Bono, 2012; Fellows & Heed, 1972).

Many members of the cactophilic *Drosophila repleta* species group can be reared under laboratory conditions without their host plant (Throckmorton, 1975; Markow & O’Grady 2008). However, *Drosophila mettleri* is an exception; this species requires its host plant to be incorporated into the food medium. One potential explanation is that either the cactus material (Fogleman et al., 1982; Markow et al., 1999; Fogleman & Danielson, 2001) or microbes present on the cactus (Starmer et al., 1976; Fogleman, Heed, et al., 1982; Fogleman, Starmer, et al., 1982; Starmer et al., 1982) provide an essential nutritional benefit to *D. mettleri*. Recently, several studies have suggested that Saguaro may provide an olfactory or gustatory cue that stimulates feeding and/or oviposition, leading to a successful laboratory culture (Benton, 2022; Álvarez-Ocaña et al., 2023). Previous studies using other cactophilic *Drosophila* species suggest that development is correlated to nutrient availability, with a preference for media containing added cactus (Markow et al., 1999; Kwang et al., 2008; Yang et al., 2008; Matzkin et al., 2011, 2013; Lihoreau et al., 2016). The process of host plant specialization can be studied by examining how Saguaro and the microbes within the cacti impact the fitness of *Drosophila mettleri* (Kibota & Courtney, 1991; Bouzas et al., 2021).

This study explores the behavioral and physiological impacts of diet in *Drosophila mettleri* and uses these results to infer long-term fitness implications of specialization to Saguaro cactus (Kristensen et al., 2003, 2016; Davies et al., 2021). We reared *D. mettleri* adults individually on cornmeal and banana media to test whether experimental trials with added Saguaro would improve *D. mettleri* pupation survival compared to control diets lacking Saguaro. We further separated the effects of nutrition from the diet or microbes within the media by repeating the experiment with microbially active Saguaro exudate and soil. We additionally considered a potential interaction with natural microbes, where the diet might either mitigate any negative effects of microbes or where the microorganisms could enhance nutrient availability from the diet (Fogleman & Danielson, 2001; Erkosar et al., 2018; Lesperance & Broderick, 2020; Davies et al., 2021; Darby et al., 2023). While we observed increased pupation survival with the addition of Saguaro, we saw little impact of microbial diversity on *D. mettleri* success. These findings highlight how the nutritional aspects of the host plant, rather than the microbes found on the necrotic tissue, directly impact *D. mettleri’s* fitness.

## Methods

### *Drosophila* and Field Sample Collection

A laboratory culture of *Drosophila mettleri* (15081-1502.15) was obtained from the National *Drosophila* Species Stock Center (NDSSC). This line was collected in La Paz, Baja California Sur, Mexico in 2012 and has been maintained on Banana media and Saguaro powder since that time (Fogelman, 2000). *Drosophila mettleri* was used for Nutritional Assays and Excrement Quantification experiments. Powdered Saguaro was maintained at -4°C in 50mL Falcon tubes. Samples were sourced from soaked soil and actively rotting Saguaro near Globe, Arizona, USA (33.57635°N, 110.95942°W). These were scraped into snap caps for Cactus Plates and Nutritional Assays.

### Nutritional Assay

The Nutritional Assay was conducted to directly compare cactus additives of varying microbial activity. We tested eight diets using either Banana or Cornmeal media as a base with one of four additive treatments: a negative control where no additives are added, and three additives (Saguaro powder from the NDSSC, Saguaro exudate from Arizona, and soil soaked with Saguaro necrosis from Arizona) (Fig 1). The banana media (Banana-Opuntia, 2018) consisted of 6% (w/v) sugar, 11% (w/v) protein, and 16% (w/v) carbohydrate. The cornmeal media (Cornell Cornmeal, 2023) consists of 4% (w/v) sugar, 9% (w/v) protein, and 15% (w/v) carbohydrate. The *Drosophila* Dietary Composition Calculator (DDCC) (Lesperance & Broderick, 2020) was developed based on previously published *Drosophila* media recipes and used to determine the exact nutrient profiles of our recipes. Each treatment had five replicates with five mating pairs in each vial (400 adult flies in total). Male and female adult flies were separately aged for 48 hours on specific treatments, then combined into vials based on their assigned treatment. Flies were held in experimental vials for 24 hours. After this time, they were serially transferred, and any eggs laid within 24 hours were counted.

**Fig 1.**
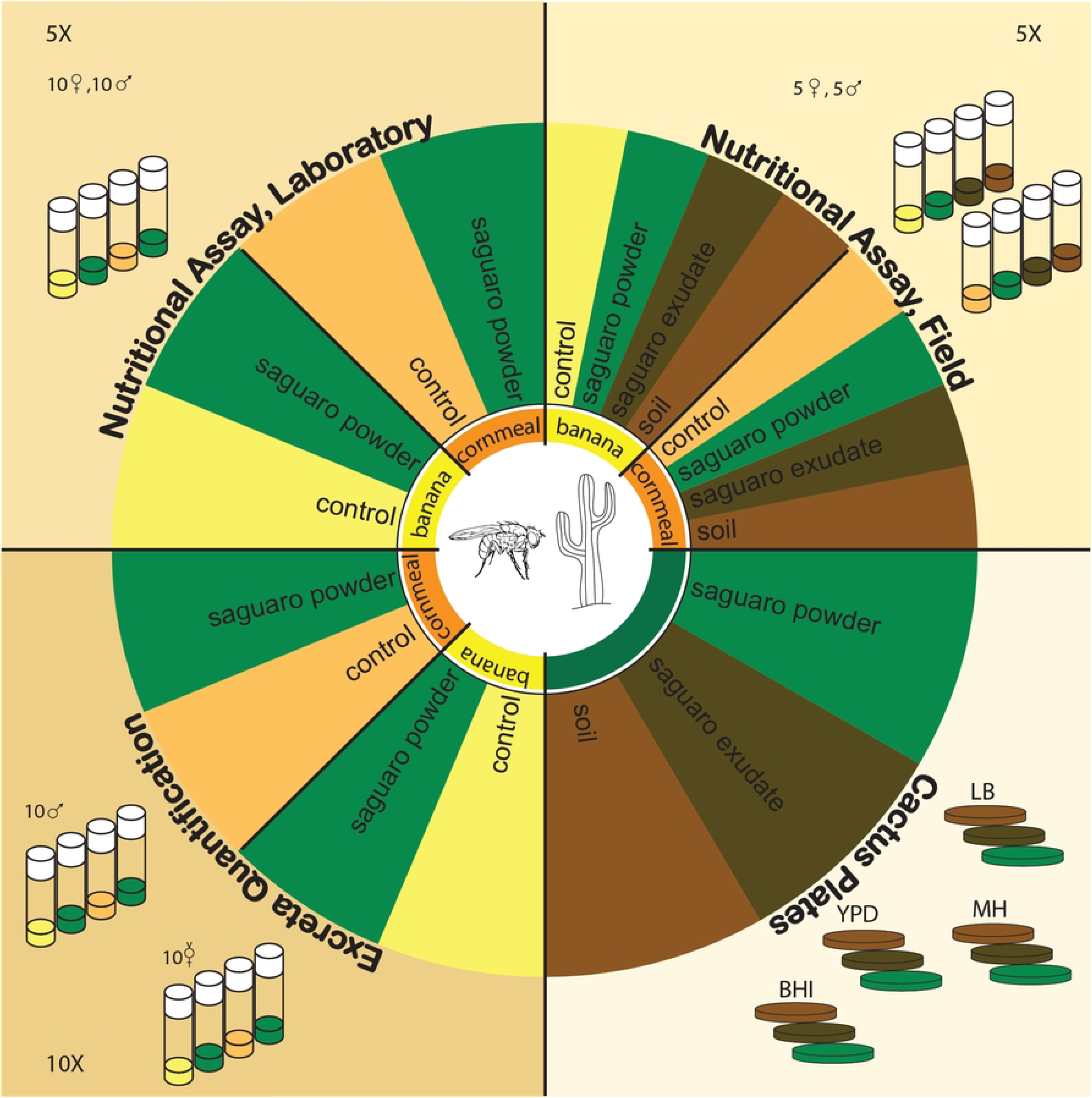
Summary of experimental layout and methods of nutritional and behavioral assays.

Adult transfer was repeated daily for five days, after which the adults were preserved in ethanol. Eggs were counted with a clicker counter and by marking the starting point, counting eggs in sight while turning clockwise. After reaching the starting point, eggs in the middle are counted from left to right using obvious markers to avoid overcounting eggs. Dates of the first larvae and first pupae per vial were noted. Larval stages could not be accurately distinguished in this rearing setup, so they were not recorded. Pupae were counted with a manual counter and marked with a colored pen every 48 hours, using a different color to denote new pupae each day of observation. Once adults emerged, they were collected every 48 hours and counted. The numbers of eggs, pupae, and adults were used to assess survival success from egg to pupa, and from pupa to adult. Survival to the pupal stage was determined by dividing the number of pupae formed by the number of eggs counted (Winkler et. al, 2021, de Paiva Mendonça et. al, 2023). The results are represented as “Pupae” (Fig 2). Pupation survival was determined by dividing the number of adults collected by the number of pupae counted (Welbergen & Sokołowski, 1994, Stobdan et. al 2024). The results are represented as “Adult” (Fig 2). Percentages were used to relative quantify survival rates for each stage. Normality of residuals was assessed using the Shapiro-Wilk test (González-Estrada, Villaseñor, & Acosta-Pech, 2022). A Kruskal-Wallis rank sum test and a Dunn’s post hoc test were conducted to examine the relationship between treatments compared to respective controls in R (Cleophas & Zwinderman 2016; Dinno & Dinno, 2017; Okoye & Hosseini, 2024). A smaller assay was also conducted with 4 treatments: Banana media with versus without cactus powder, versus Cornmeal media with versus without cactus powder. This assay serves to measure the baseline developmental rate of *Drosophila mettleri* on Banana and Cornmeal diet (Table S3).

**Fig 2.**
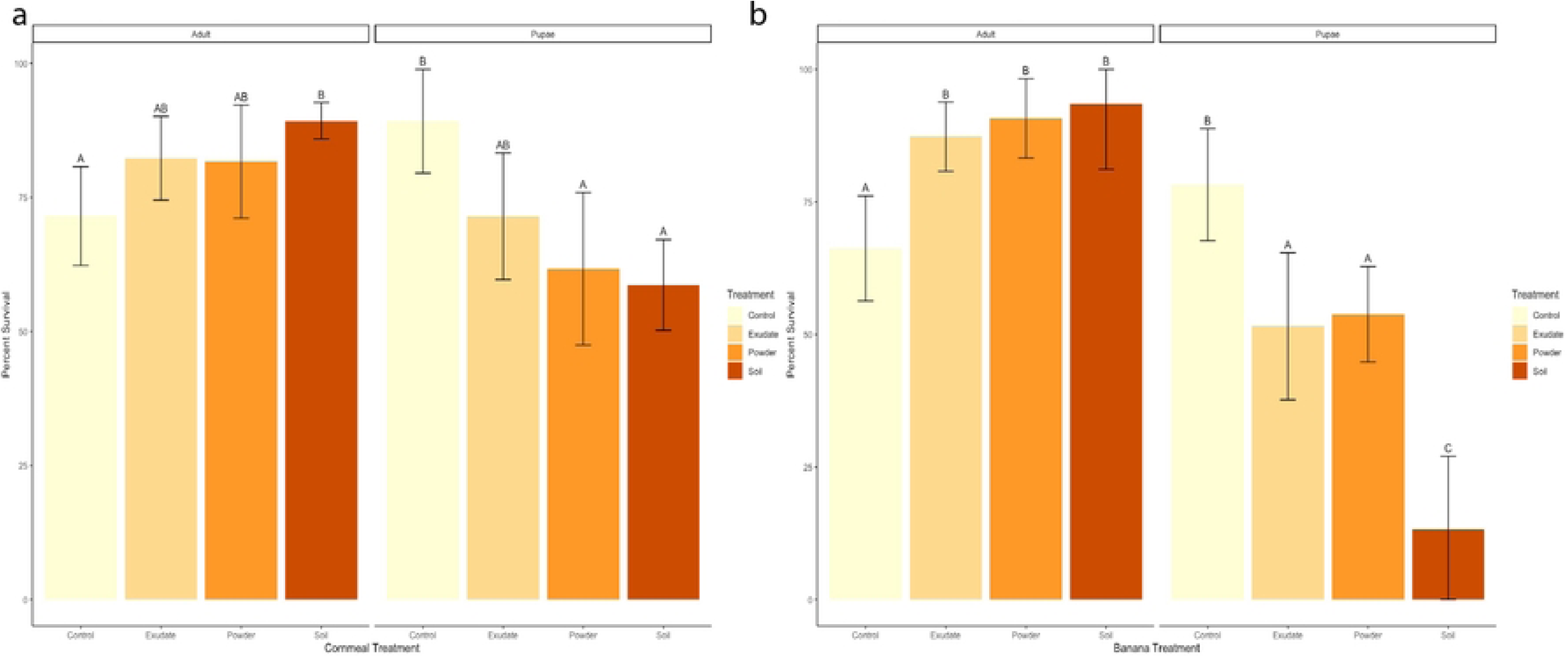
Nutritional Assay Results. Mean (± 95% C.I.) Pupae and Adult survival rates of *D. mettleri* on control (yellow), Saguaro exudate (tan), Saguaro powder (orange), and Saguaro soil (brown) on (a) Cornmeal and (b) Banana media. Letters above error bars indicate difference between the means, with identical letters indicating results are not significantly different based on Dunn’s post hoc test (p < 0.05).

### Excreta Quantification Assay

The Excreta Quantification (Ex-Q) Assay is adapted from Wu et. al (2020), and is used to assess whether adults fed differently on media with cactus present. The assay was conducted with four treatments: Banana media with and without cactus powder, and Cornmeal with and without cactus powder (Fig 1). Male and unmated female adult flies were evaluated in separate trials and aged separately on their treatments for three days before they were used for the experiment. Ex-Q tubes were created from 50 mL conical tubes with a hole cut on the top where a 1.5 mL centrifuge tube lid was placed and used as a food cup. The food cup was filled with 50 uL of dyed treatment food. Food was colored by mixing powder of the non-digestible food dye erioglaucine 1% (w/v). Each treatment had 5 replicates with 10 individuals in each vial. Flies were serially transferred to a new Ex-Q tube every 24 hrs for 2 days, then transferred to an empty vial for 3 hours to excrete the remaining dye. Excreted dye was collected by adding 3 mL of deionized water to the Ex-Q tube, taping the top opening, and spinning down the contents of the tube. Absorbance was measured at 630 nm in a 96-well plate on a spectrometer (Molecular Biosciences Spectra Max Series Plus 384). Each plate contained samples from all eight treatments over the total number of days. A standard curve was prepared by dissolving 10 mg of erioglaucine in 10 mL of PBS followed by a serial 2-fold dilution. The final range of standards was from 78 μg/mL to 5 μg/mL. A 100 uL aliquot of standard (2 replicates) and treatments (3 replicate; 3 flies per treatment) were dispensed into individual wells of a 96-well plate. Each repeat was averaged for the final data analysis. The spectrophotometer absorbance readings were used to calculate the average amount of dye the flies excreted after feeding on the various diets. The collected dye therefore approximates how much the flies ate on each treatment based on the following equation: Dye (μg) per fly = [(Absorbance–Intercept)*dye collection volume]/(Slope * # flies*aliquot volume). Normality of residuals was assessed using the Shapiro-Wilk test (González-Estrada, Villaseñor, & Acosta-Pech, 2022). A Kruskal-Wallis rank sum test was conducted in R to examine diet consumption of male and female flies provided with media that contained cactus powder compared to controls for each food type (Dinno & Dinno, 2017; Okoye & Hosseini, 2024).

### Cactus plates

Aliquots of Saguaro powder, exudate, and soil were cultured and plated to assess microbial community. Four types of microbial growth plates were used for this experiment: Yeast Peptone Dextrose (YPD), Luria Broth (LB) Agar, Brain Heart Infusion (BHI), and Mueller Hinton (MH). Three types of cactus samples were used: one control powder and two field samples (Fig 1). The Saguaro powder used for the Nutritional Assay was frozen, defrosted, and opened multiple times. The two field samples were exudate and soil that was soaked in cactus exudate collected from Globe, Arizona. The three types of Saguaro will henceforth be called “treatments.” Deionized water was used as a negative control. To plate, 14 mg of each treatment was suspended in 1 mL of deionized water and centrifuged for 10 seconds. The mixture was resuspended by pipetting up at down at least three times immediately prior to plating. A 40 uL aliquot of the suspension was plated on each type of plate (equating to about 56ug of cactus added to each plate) and spread with glass beads. The plates were incubated at 21°C, which is the temperature at which the experiments using live *D. mettleri* were conducted. After 24 hours, images of each plate were taken. Plate pictures were processed using a plate count protocol in ImageJ to assess the number of colonies grown on each medium (Jeric 2016). Data were analyzed using a two-way analysis of variance (ANOVA) to assess the effects of cactus and plate type on the number, size, and total area of microbial colonies (Okoye & Hosseini, 2024).

Significance was determined based on the p-values, with Tukey’s Honest Significant Difference (HSD) test used for post-hoc pairwise comparisons (Abdi & Williams 2010). Normality of residuals was assessed using the Shapiro-Wilk test (González-Estrada, Villaseñor, & Acosta-Pech, 2022).

## Results

### Ex-Q Assay

Adults on Cornmeal showed a slight increase in the amount of dye excreted compared to adults on Banana media, but these results were not significant by a Kruskal-Wallis rank sum test (Fig 2a, 2b). Unmated males and females on food with cactus showed no significant difference in the amount of dye excreted when compared to respective controls (Fig 3a, 3b). This indicates that the flies eat the same amount on each diet treatment, and any developmental differences did not arise due to consumption differences.

**Fig 3.**
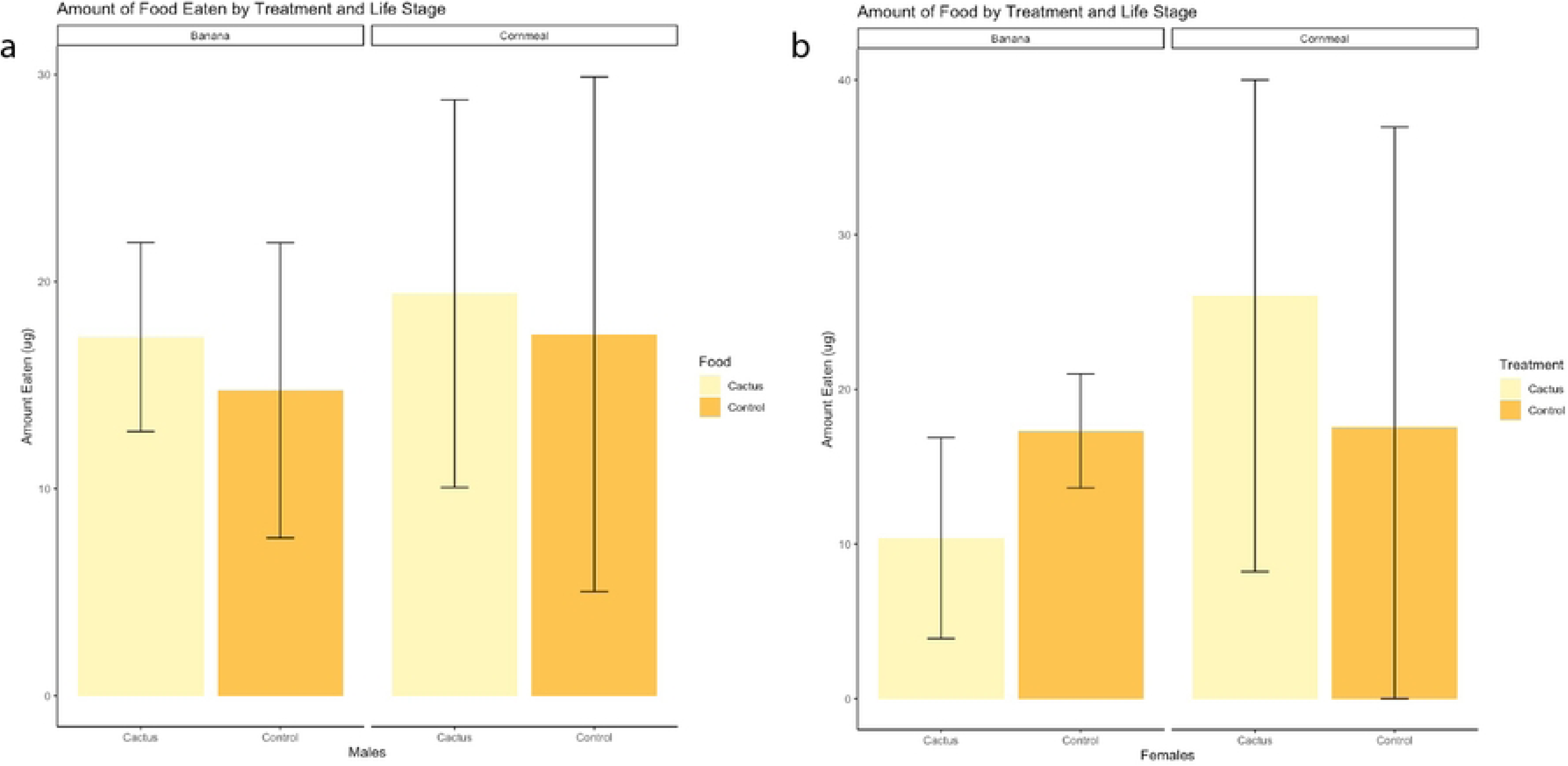
Excrement Quantitification Assay Results. Mean (± 95% C.I.) amount of dye excreted in ug by males (a) and unmated females (b) on Banana and Cornmeal with Saguaro powder added and controls. Bars without letters are not significantly different based on Dunn’s post hoc test (p < 0.05).

### Nutritional Assay

*Drosophila mettleri* survival differed across media type, cactus treatment, and life stage assessed (Fig 2, Fig S3). Survival to the pupal stage was lower in all three cactus treatments than in controls (13-54% vs. 78% in Banana, 58-71% vs 89% in Cornmeal; Fig 2a, 2b, 5, Fig S3a, S3b, S5). However, the percentage of individuals surviving pupation and emerging as adults was significantly higher in cactus treatments than controls (87-93% vs 66% in Banana, 81-89% vs 71% in Cornmeal; Fig 2a, 2b, 5, Fig S3a, S3b, S5). Interestingly, there were no significant survival differences between most types of cactus additives on Banana or Cornmeal, except that pupal survival was dramatically worse on the Banana-soil combination (Fig 2a, 2b, 5, Fig S3a, S3b, S5).

The duration from the first larvae to pupae and from pupae to adults was not significantly different across media (Table S1). However, *Drosophila* reared on media containing Saguaro soil consistently took 48hrs longer to develop, while Drosophila reared on Saguaro exudate or powder developed 24-48hrs shorter compared to controls (Table S1).

### Cactus Plates

Saguaro powder, exudate, and soaked soil were plated on a variety of microbial culture media to determine whether viable yeast and bacteria were present (Fig 4). The Saguaro powder showed the lowest level of microbial activity, with either no or low (0-6.25 colonies) culture growth (Fig 4, Table S2). In contrast, Saguaro exudate and soil samples had more colonies (987- 1710 colonies) and total area of growth (4164-4185 mm^2^ vs 216 mm^2^), implying higher microbial activity than plates with powder samples (Fig 3, Table S2). While plates with Saguaro soil had colony sizes twice as large as exudate samples, Saguaro exudate had almost twice as many microbial colonies potentially indicating different microbial community between the two additives or that Saguaro exudate has higher microbial activity relative to Saguaro soil (Fig 4, Table S2).

**Fig 4.**
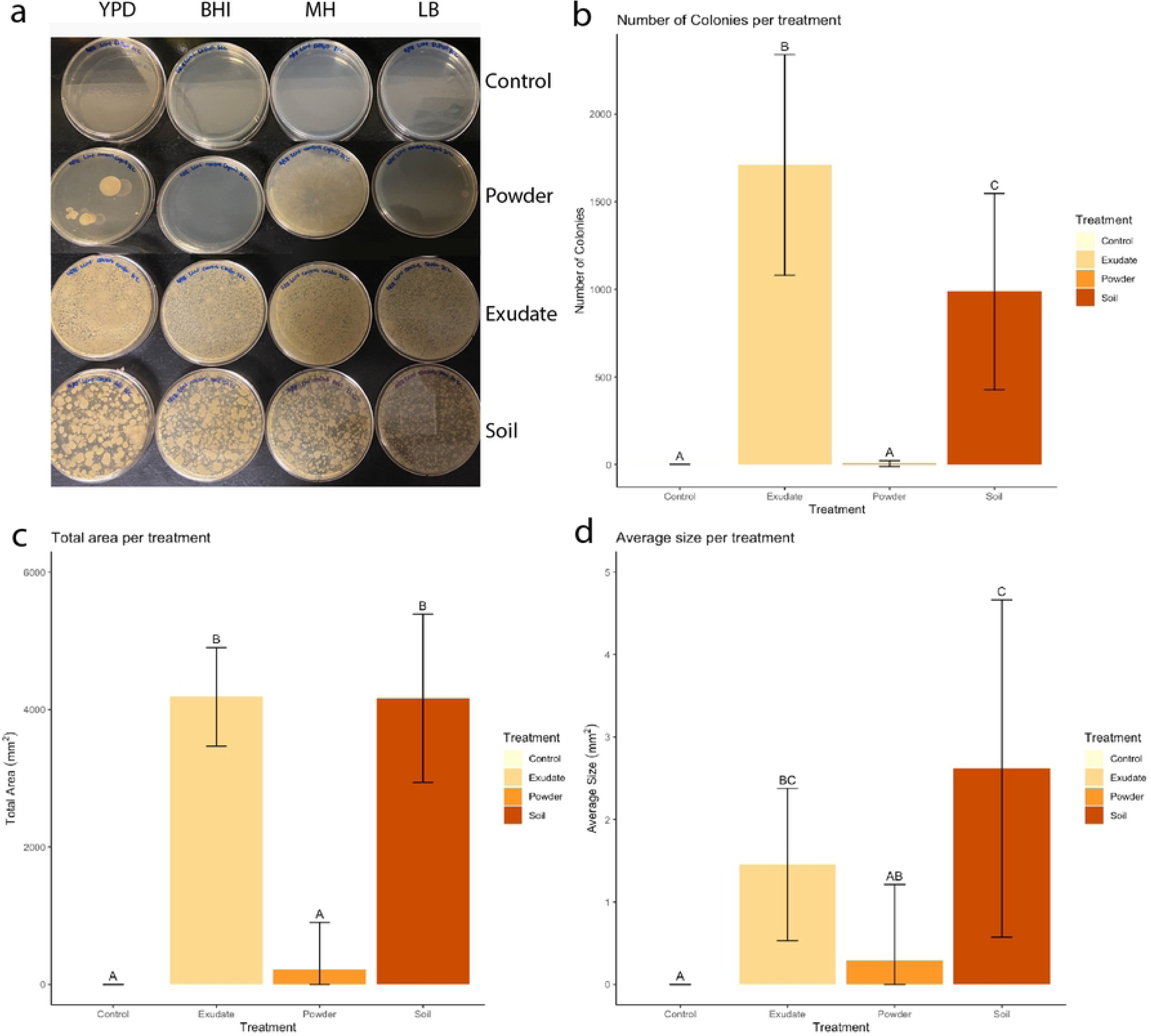
Cactus Plates Results. (a) YPD, BHI, MH, and LB plates depicting microbial growth of Control, Powder, Exudate, and Soil samples. Graphs of (b) Total Number of Colonies, (c) Total Colony Area, and (d) Average Colony Size per treatment of control (yellow), Saguaro exudate (tan), Saguaro powder (brown). Letters above error bars indicate difference between the means, with identical letters showing not significantly different based on Tukey’s HSD post hoc test (p < 0.05).

**Fig 5.**
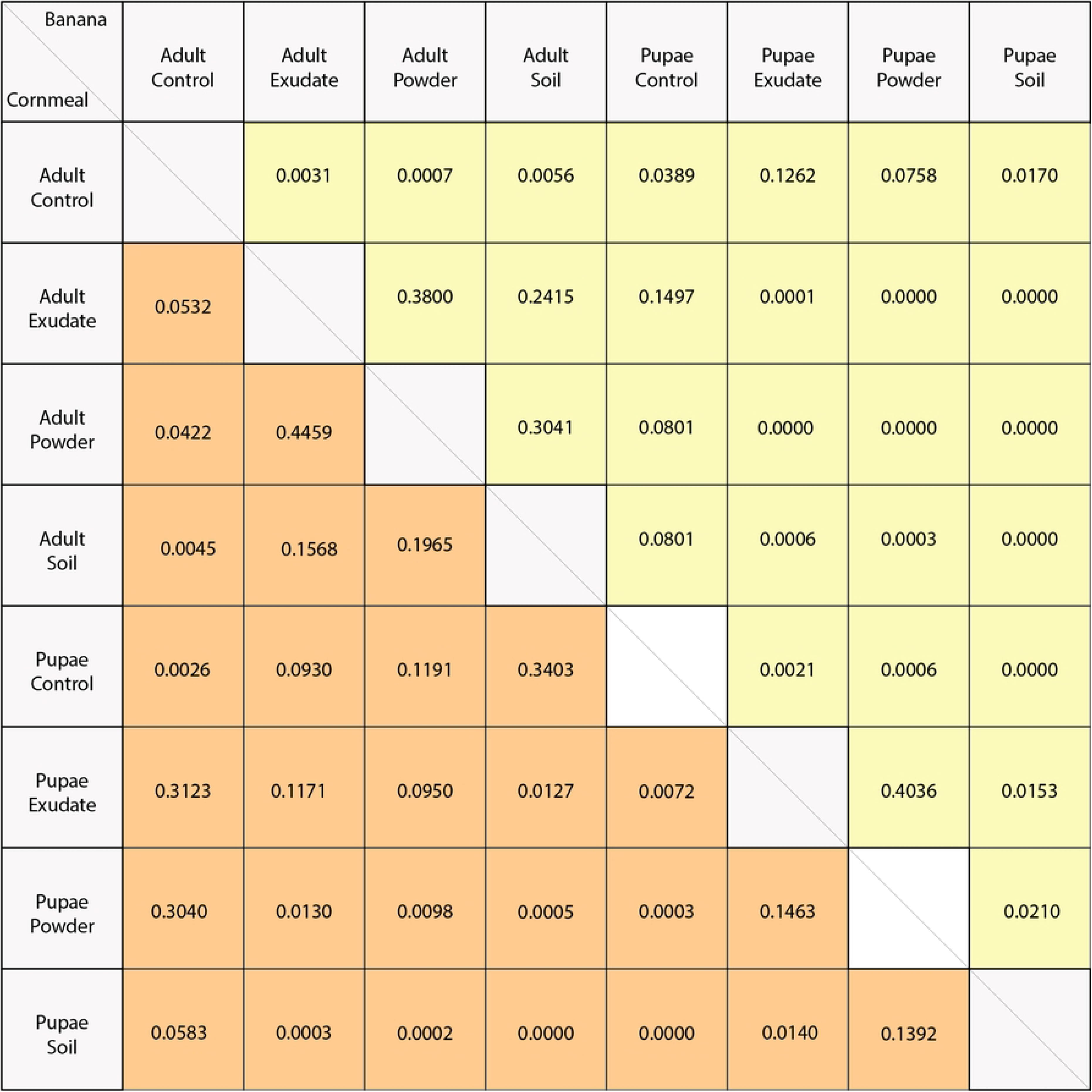
Nutritional Assay Dunn’s post hoc test results. Banana (yellow) and Cornmeal (orange) treatments in Nutritional Assay.

## Discussion

In this study, we evaluated how various Saguaro preparations and media types affect *D. mettleri* pupation success. We found higher pupation survival after supplementation with cactus for cornmeal and banana media, with a stronger response in the banana treatment (Fig 2a, 2b, Fig S3a, S3b) despite equivalent *D. mettleri* consumption across the diets (Fig 3a, 3b). This suggests that an increase in bioavailable nutrients from the cactus provides a developmental benefit. These results are in line with other studies on specialist *Drosophila* species, where *mojavensis*, *elegans*, and *sechellia* larvae had lower body weights, higher mortality, and delayed development when compared to *melanogaster* and/or *simulans* on high sugar, lower protein diets (Matzkin et al., 2011; Watanabe et al., 2019).

Pre-pupation success can be influenced by the environment of the fly, including by larval competition (Matzkin et al., 2013; Vijendravarma et al., 2013; Ahmad et al., 2015; Hardin et al., 2015). In the wild, the adults lay hundreds of eggs on a suitable larval development substrate.

Once the eggs hatch, the hundreds of larvae begin to compete with one another for resources. Larvae with an earlier hatching date gain precedence on resources within the environment, influencing success in the later stages. While the larvae have more substrate to explore in the wild, they are confined to a certain amount of food and space within a tube under lab conditions. The finite resources lead to heightened larval competition, resulting in potential delayed development, reduced size, or increased mortality (Vijendravarma et al., 2013; Ahmad et al., 2015; Hardin et al., 2015). The addition of powdered cactus and microbially active exudate adds another layer of complexity by changing the environment. The cactus boosts the nutritional value of the media by providing sterols and simple chemicals that lab media generally lacks (Fogleman, Heed, et al., 1982; Fogleman & Danielson, 2001). These additions can promote faster growth and shorter development times, which explains the development patterns observed in flies reared on Saguaro powder and exudate (Table S1) (Kwang et al., 2008; Lihoreau et al., 2016; Matzkin et al., 2011; Yang et al., 2008).

Depending on the interplay between a poor diet and heightened larval competition, improving the nutritional quality of the media may not prove beneficial (Hardin et al., 2015). Supplementation of the laboratory diet with microbially active soil from the natural *D. mettleri* environment. Some soil microbes can enhance nutrient availability by either breaking down cactus into more bioavailable constituents, or the microbes themselves may serve as a nutrient source for the *Drosophila* (Altomare et al., 2011; Miransari, 2013; Sahu et al., 2018; Lesperance & Broderick, 2020; Davies et al., 2021). At the same time, some of these microbes can consume nutrients in the substrate and may proliferate faster than larvae can develop, such that the microbes overtake the environment and compete with larvae for resources (Vijendravarma et al., 2013; Ahmad et al., 2015). These factors have fitness consequences for individual flies and may collectively influence pre-pupation success rates.

The Cactus Plates assay (Fig 4) explores the microbial diversity in the different treatments. Cactus powder had the lowest microbial activity, Saguaro exudate had the highest, and Saguaro soil was intermediate in activity (Table S2). Lower microbial activity in the cactus powder is expected since it underwent extreme conditions during its manufacturing and storage, probably limiting the microbial community that could survive. While a large component of Saguaro soil treatments was Saguaro exudate, the number of recovered microbial colonies was about half in soil relative to exudate. This suggests that not all microbes present in cactus exudate survive in the soil. However, we observed that soil microbes improve *D. mettleri* developmental outcomes. Significant adult *D. mettleri* survival in soil treatments replicated over both media types further demonstrates that Saguaro soil is a better additive compared to powder and exudate for developmental success. The observed variability in adult success rates across cactus additives indicates that factors beyond substrate type may influence the later stages of *Drosophila* development. These factors could include nutritional content, microbial activity, or other environmental conditions.

The amount of media consumed by flies on the Cornmeal diet compared to the Banana diet did not differ for unmated females or males and is not in line with the results of similar studies (Wu et. al, 2020). According to the *Drosophila* Dietary Composition Calculator (DDCC), the Cornmeal diet with no cactus additions has a protein: carbohydrate ratio (P:C) of 1:3.5 (Lesperance & Broderick, 2020). The Banana diet with no cactus additions has a protein: carbohydrate ratio of 1:11, indicating this diet has 3 times more carbohydrates than the Cornmeal diet. We added 14mg/100g of cactus powder, which adds 4.69mg/100g of protein and 1.44mg/100g of carbohydrates, significantly increasing the protein: carbohydrate ratios to 1:3 and 1:8 for Cornmeal and Banana, respectively (Shalaby, Ghanem, & Maamon, 2016). Adult *Drosophila melanogaster* fed on a high sugar diet (> 8% w/v) consume a significantly lower amount than those fed on a low sugar diet (< 4% w/v) (van Dam et al., 2020; Darby et al., 2023). Our present data suggest that, unlike *D. melanogaster*, *D. mettleri* do not exhibit feeding differences based on the amount of sugar (or other Saguaro macronutrients) in the diet. This may reflect dramatic differences in the natural ecology of the two flies, as *D. melanogaster* feeds and develops on sugar-rich fermenting fruit in nature (Throckmorton, 1975; Markow and O’Grady, 2005).

Our experimental design did not allow for interaction between unmated females and males. The results we observe may have been biased because females in the absence of males may have more time to feed, or because mated females may increase feeding relative to unmated females because of the nutritional requirements of egg production. Mated *Drosophila melanogaster* females are more likely to make nutritional choices based on the ovipositional site that will provide their progeny with the best fitness outcomes (Yang et al., 2008; Lee et al., 2013; Matzkin et al., 2013; Lihoreau et al., 2016). In *D. melanogaster*, unmated females and males tend to consume more carbohydrates in their diet (P:C 1:4) while mated females tend to consume comparatively more protein (P:C 1:1.5) (Kwang et al., 2008; Lee et al., 2013). While *D. melanogaster* and *D. mettleri* are taxonomically distant, their nutritional requirements could be evolutionarily conserved as a function of the need to provision eggs.

Our results demonstrate that *Drosophila mettleri* derive direct fitness benefits from its specialized host plant, *Carnegiea gigantea*, supporting the idea that nutrition plays a central role in maintaining ecological specialization. While microbial communities may influence development under certain conditions, our findings indicate that it is the nutritional properties of the host plant itself that are most critical for survival and successful pupation. In a resource-limited environment of the Sonoran Desert, such a dependency can be a competitive advantage by enabling *D. mettleri* to exploit a niche that is inaccessible to other organisms. More broadly, our study highlights how host plant chemistry can directly shape fitness in specialist insects, offering insight into the mechanisms underlying host-use evolution and the persistence of specialization under extreme environmental pressures.. These results add to the growing field of nutritional ecology and lay the groundwork for understanding physiological processes leading to *D. mettleri’s* adaptation to Saguaro cactus.

## Conclusions

While all cactus additives improved pupation success, Saguaro soil consistently yielded the highest adult survival, likely due to a combination of increased nutrient availability and microbial contributions. However, feeding rates remained unchanged across diets, suggesting that the observed developmental differences arise not from intake quantity but from the nutritional or microbial quality of the substrate. This supports the idea that *D. mettleri*’s specialization on *Carnegiea gigantea* may reflect a dependency shaped by ecological pressures, though whether these responses are adaptive or by-products of environmental constraints requires further investigation.

## Acknowledgments

We thank the Saguaro National Parks (permit number SAGU-2023-SCI-0007) and park rangers for their support throughout the study. We thank Benjamin Burgunder, Andrew Legan, Alan Mata, Kyla O’Hearn, and Augusto Santos Rampasso for their assistance in the field.

## Supporting information

**Table S1. Table of mean survival and development times (days).** Table shows egg, pupa, and adult *D. mettleri* results for the Nutritional Assay showed in Fig 1.

**Table S2. Table of agar plate readings**. Count indicates the number of colonies found. Total area refers to the total area of microbial coverage. Average size refers to the average size of colonies. %Area indicates the percentage colony cover compared to the total area. Mean refers to the average pixel value.

**Fig S3. Supplementary Nutritional Assay Results**. Mean (± 95% C.I.) Pupae and Adult survival rates of *D. mettleri* on control (yellow) and Saguaro powder (orange) on Banana (a) and Cornmeal (b) media. Letters above error bars indicate difference between the means with identical letters showing not significantly different based on Dunn’s post hoc test (p < 0.05).

**Table S4. Table of survival percentages and development times (days).** Table shows egg, pupa, and adult *D. mettleri* results for the additional Nutritional Assay shown in Fig S3.]

**Fig S5. Supplementary Nutritional Assay Dunn’s post hoc test results.** Banana (yellow) and Cornmeal (orange) for the additional Nutritional Assay in Fig S3a and S3b.

## References

Abdi, H., & Williams, L. J. (2010). Tukey’s honestly significant difference (HSD) test. Encyclopedia of research design, 3(1), 1–5.

Ahmad, M., Chaudhary, S. U., Afzal, A. J., & Tariq, M. (2015). Starvation-Induced Dietary Behaviour in Drosophila melanogaster Larvae and Adults. Scientific Reports 2015 5:1, 5(1), 1–13. 10.1038/srep14285

Altomare, C., Tringovska, I., Altomare, C., Tringovska Maritsa, I., & Lichtfouse, E. (2011). Beneficial Soil Microorganisms, an Ecological Alternative for Soil Fertility Management. 7, 161–214. 10.1007/978-94-007-1521-9_6

Álvarez-Ocaña, R., Shahandeh, M. P., Ray, V., Auer, T. O., Gompel, N., & Benton, R. (2023). Odor-regulated oviposition behavior in an ecological specialist. Nature Communications, 14(1), 3041. 10.1038/s41467-023-38722-z

Banana-Opuntia. National Drosophila Species Stock Center. (2018, July 9). https://www.drosophilaspecies.com/recipes/banana-opuntia/

Benton, R. (2022). Drosophila olfaction: Past, present and future. Proceedings of the Royal Society B: Biological Sciences, 289(1989). 10.1098/rspb.2022.2054

Bouzas, S., Barbarich, M. F., Soto, E. M., Padró, J., Carreira, V. P., & Soto, I. M. (2021). Specialization and performance trade-offs across hosts in cactophilic Drosophila species. Ecological Entomology, 46(4), 877–888. 10.1111/een.13024

Castrezana, S., & Bono, J. M. (2012). Host Plant Adaptation in Drosophila mettleri Populations. PLOS ONE, 7(4), e34008. 10.1371/journal.pone.0034008

Coleman, D. C., Crossley, D. A., & Hendrix, P. F. (2004). Secondary Production: Activities of Heterotrophic Organisms—The Soil Fauna. Fundamentals of Soil Ecology, 79–185. 10.1016/B978-012179726-3/50005-8

Cornell cornmea*l*. National Drosophila Species Stock Center. (2023, December 26). https://www.drosophilaspecies.com/recipes/cornell-cornmeal/

Cleophas, T.J., Zwinderman, A.H. (2016). Non-parametric Tests for Three or More Samples (Friedman and Kruskal-Wallis). In: Clinical Data Analysis on a Pocket Calculator. Springer, Cham. 10.1007/978-3-319-27104-0_34

Darby, A. M., Okoro, D. O., Aredas, S., Frank, A. M., Pearson, W. H., Dionne, M. S., & Lazzaro, B. P. (2023). High sugar diets can increase susceptibility to bacterial infection in Drosophila melanogaster. BioRxiv, 2023.12.07.570705. 10.1101/2023.12.07.570705

Dinno, A., & Dinno, M. A. (2017). Package ‘dunn. test’. CRAN Repos, 10, 1–7.

Davies, L. R., Loeschcke, V., Schou, M. F., Schramm, A., & Kristensen, T. N. (2021). The importance of environmental microbes for Drosophila melanogaster during seasonal macronutrient variability. Scientific Reports 2021 11:1, 11(1), 1–11. 10.1038/s41598-021-98119-0

Duek, L., Kaufman, G., Palevsky, E., & Berdicevsky, I. (2001). Mites in fungal cultures. Mycoses, 44(9–10), 390–394. 10.1046/j.1439-0507.2001.00684.x

Erkosar, B., Yashiro, E., Zajitschek, F., Friberg, U., Maklakov, A. A., van der Meer, J. R., & Kawecki, T. J. (2018). Host diet mediates a negative relationship between abundance and diversity of Drosophila gut microbiota. Ecology and Evolution, 8(18), 9491–9502. 10.1002/ECE3.4444

Fellows, D. P., & Heed, W. B. (1972). Factors Affecting Host Plant Selection in Desert-Adapted Cactiphilic Drosophila. Ecology, 53(5), 850–858. 10.2307/1934300

Fogleman, J. C. (2000). Response of Drosophila melanogaster to selection for P450- mediated resistance to isoquinoline alkaloids, Chemico-Biological Interactions, 125 (2), 93–105.10.1016/S0009-2797(99)00161-1.

Fogleman, J. C., & Danielson, P. B. (2001). Chemical Interactions in the Cactus- Microorganism-Drosophila Model System of the Sonoran Desert1. American Zoologist, 41(4), 877–889. 10.1093/icb/41.4.877

Fogleman, J. C., Heed, W. B., & Kircher, H. W. (1982). Drosophila mettleri and senita cactus alkaloids: Fitness measurements and their ecological significance. Comparative Biochemistry and Physiology Part A: Physiology, 71(3), 413–417. 10.1016/0300-9629(82)90427-3

Fogleman, J. C., Starmer, W. T., & Heed, W. B. (1982). Comparisons of yeast florae from natural substrates and larval guts of southwestern Drosophila. Oecologia, 52(2), 187–191. 10.1007/BF00363835/METRICS

González-Estrada, E., Villaseñor, J. A., & Acosta-Pech, R. (2022). Shapiro-Wilk test for multivariate skew-normality. Computational Statistics, 37(4), 1985–2001.

Hardin, J. A., Kraus, D. A., & Burrack, H. J. (2015). Diet quality mitigates intraspecific larval competition in Drosophila suzukii. Entomologia Experimentalis et Applicata, 156(1), 59–65. 10.1111/EEA.12311

Kibota, T. T., & Courtney, S. P. (1991). Jack of one trade, master of none: host choice by Drosophila magnaquinaria. Oecologia, 86(2), 251–260. 10.1007/BF00317538/METRICS

Kristensen, T. N., Dahlgaard, J., & Loeschcke, V. (2003). Effects of inbreeding and environmental stress on fitness – using Drosophila buzzatii as a model organism. Conservation Genetics, 4(4), 453–465. 10.1023/A:1024763013798

Kristensen, T. N., Henningsen, A. K., Aastrup, C., Bech-Hansen, M., Bjerre, L. B. H., Carlsen, B., Hagstrup, M., Jensen, S. G., Karlsen, P., Kristensen, L., Lundsgaard, C., Møller, T., Nielsen, L. D., Starcke, C., Sørensen, C. R., & Schou, M. F. (2016). Fitness components of Drosophila melanogaster developed on a standard laboratory diet or a typical natural food source. Insect Science, 23(5), 771–779. 10.1111/1744-7917.12239

Kwang, P. L., Simpson, S. J., Clissold, F. J., Brooks, R., Ballard, J. W. O., Taylor, P. W., Soran, N., & Raubenheimer, D. (2008). Lifespan and reproduction in Drosophila: New insights from nutritional geometry. Proceedings of the National Academy of Sciences of the United States of America, 105(7), 2498–2503. 10.1073/PNAS.0710787105/SUPPL_FILE/LEE_ET_AL._SUPPORTING_INFO_PNAS_2008.PDF

Lee, K. P., Kim, J. S., & Min, K. J. (2013). Sexual dimorphism in nutrient intake and life span is mediated by mating in Drosophila melanogaster. Animal Behaviour, 86(5), 987– 992. 10.1016/J.ANBEHAV.2013.08.018

Lesperance, D. N. A., & Broderick, N. A. (2020). Meta-analysis of Diets Used in Drosophila Microbiome Research and Introduction of the Drosophila Dietary Composition Calculator (DDCC). G3 Genes|Genomes|Genetics, 10(7), 2207–2211. 10.1534/G3.120.401235

Lihoreau, M., Poissonnier, L. A., Isabel, G., & Dussutour, A. (2016). Drosophila females trade off good nutrition with high-quality oviposition sites when choosing foods. Journal of Experimental Biology, 219(16), 2514–2524. 10.1242/JEB.142257/262455/AM/DROSOPHILA-FEMALES-TRADE-OFF-GOOD-NUTRITION-WITH

Markow, T. A., & O’Grady, P. (2008). Reproductive ecology of Drosophila. Functional Ecology, 22(5), 747–759. 10.1111/j.1365-2435.2008.01457.x

Markow, T. A., Raphael, B., Dobberfuhl, D., Breitmeyer, C. M., Elser, J. J., & Pfeiler, E. (1999). Elemental stoichiometry of Drosophila and their hosts. Functional Ecology, 13(1), 78–84. 10.1046/J.1365-2435.1999.00285.X

Matzkin, L. M., Johnson, S., Paight, C., Bozinovic, G., & Markow, T. A. (2011). Dietary Protein and Sugar Differentially Affect Development and Metabolic Pools in Ecologically Diverse Drosophila. The Journal of Nutrition, 141(6), 1127–1133. 10.3945/JN.111.138438

Matzkin, L. M., Johnson, S., Paight, C., & Markow, T. A. (2013). Preadult Parental Diet Affects Offspring Development and Metabolism in Drosophila melanogaster. PLOS ONE, 8(3), e59530. 10.1371/JOURNAL.PONE.0059530

Meyer, J. M., & Fogleman, J. C. (1987). Significance of Saguaro cactus alkaloids in ecology of Drosophila mettleri, a soil-breeding, cactophilic drosophilid. Journal of Chemical Ecology, 13(11), 2069–2081. 10.1007/BF01012872

Miransari, M. (2013). Soil microbes and the availability of soil nutrients. Acta Physiologiae Plantarum, 35(11), 3075–3084. 10.1007/S11738-013-1338-2/TABLES/2

Okoye, K., Hosseini, S. (2024). Analysis of Variance (ANOVA) in R: One-Way and Two-Way ANOVA. In: R Programming. Springer, Singapore. 10.1007/978-981-97-3385-9_9

Okoye, K., & Hosseini, S. (2024). R Programming: Statistical Data Analysis in Research. Springer Nature.

de Paiva Mendonça L, Haddi K, Godoy WAC (2023) Effects of co-occurrence and intra- and interspecific interactions between Drosophila suzukii and Zaprionus indianus. PLOS ONE 18(3): e0281806. 10.1371/journal.pone.0281806

Saguaro - Saguaro National Park (U.S. National Park Service). (n.d.). Retrieved April 15, 2024, from https://www.nps.gov/sagu/learn/nature/Saguaro.htm

Sahu, A., Bhattacharjya, S., Mandai, A., Thakur, J. K., Atoliya, N., Sahu, N., Manna, M. C., & Patra, A. K. (2018). Microbes: A sustainable approach for enhancing nutrient availability in agricultural soils. Role of Rhizospheric Microbes in Soil: Volume 2: Nutrient Management and Crop Improvement, 47–75. 10.1007/978-981-13-0044-8_2/TABLES/6

Sánchez-Chávez, D. I., Rodríguez-Zaragoza, S., Velez, P., Cabirol, N., & Ojeda, M. (2023). Fungal feeding preferences and molecular gut content analysis of two abundant oribatid mite species (Acari: Oribatida) under the canopy of Prosopis laevigata (Fabaceae) in a semi-arid land. Experimental & Applied Acarology, 89(3–4), 417. 10.1007/S10493-023-00790-7

Selden, P. A. (2017). Arachnids. Reference Module in Life Sciences. 10.1016/B978-0-12-809633-8.02243-3

Shalaby, M., Ghanem, A., & Maamon, H. (2016). Protective effect of ginger and cactus saguaro extract against Cancer Formation Cells. Journal of Food and Dairy Sciences, 7(11), 487–491. 10.21608/jfds.2016.46069

Sharma, G., Malthankar, P. A., & Mathur, V. (2020). Insect–Plant Interactions: A Multilayered Relationship. Annals of the Entomological Society of America, 114, 1–16. https://api.semanticscholar.org/CorpusID:225144955

Starmer, W., Phaff, H. J., Miranda, M., Miller, M. W., & Heed, W. B. (1982). The yeast flora associated with the decaying stems of columnar cacti and Drosophila in North America. Evolutionary Biology, 14, 269–295.

Starmer, W. T., Heed, W. B., Miranda, M., Miller, M. W., & Phaff, H. J. (1976). The ecology of yeast flora associated with cactiphilic Drosophila and their host plants in the Sonoran desert. Microbial Ecology, 3(1), 11–30. 10.1007/BF02011450

Stobdan, T., Wen, N. J., Lu-Bo, Y., Zhou, D., & Haddad, G. G. (2024). The Pupa Stage Is the Most Sensitive to Hypoxia in Drosophila melanogaster. International Journal of Molecular Sciences, 25(2), 710. 10.3390/ijms25020710

Throckmorton, L. H., (1982). Pathways of evolution in the genus Drosophila and the founding of the repleta group, pp. 33–47 in Ecological Genetics and Evolution: the Cactus-Yeast-Drosophila Model System, edited by J. F. S. Barker and W. T. Starmer. Academic Press, New York

van Dam, E., van Leeuwen, L. A. G., dos Santos, E., James, J., Best, L., Lennicke, C., Vincent, A. J., Marinos, G., Foley, A., Buricova, M., Mokochinski, J. B., Kramer, H. B., Lieb, W., Laudes, M., Franke, A., Kaleta, C., & Cochemé, H. M (2020). Sugar-Induced Obesity and Insulin Resistance Are Uncoupled from Shortened Survival in Drosophila. Cell Metabolism, 31(4), 710. 10.1016/J.CMET.2020.02.016

Vijendravarma, R. K., Narasimha, S., & Kawecki, T. J. (2013). Predatory cannibalism in Drosophila melanogaster larvae. Nature Communications 2013 4:1, 4(1), 1–8. 10.1038/ncomms2744

Watanabe, K., Kanaoka, Y., Mizutani, S., Uchiyama, H., Yajima, S., Watada, M., Uemura, T., & Hattori, Y. (2019). Interspecies Comparative Analyses Reveal Distinct Carbohydrate-Responsive Systems among Drosophila Species. Cell Reports, 28(10), 2594–2607.e7. 10.1016/J.CELREP.2019.08.030

Welbergen, P., Sokolowski, M.B. Development time and pupation behavior in theDrosophila melanogaster subgroup (Diptera: Drosophilidae). J Insect Behav 7, 263– 277 (1994). 10.1007/BF01989734

Winkler, A., Jung, J., Kleinhenz, B. and Racca, P. (2021), Estimating temperature effects on *Drosophila suzukii* life cycle parameters. Agr Forest Entomol, 23: 361–377. 10.1111/afe.12438

Winkler, I. S., & Mitter, C. (2008). 240 The Phylogenetic Dimension of Insect-Plant Interactions: A Review of Recent Evidence. In K. Tilmon (Ed.), Specialization, Speciation, and Radiation: The Evolutionary Biology of Herbivorous Insects (p. 0). University of California Press. 10.1525/california/9780520251328.003.0018

Yang, C. H., Belawat, P., Hafen, E., Jan, L. Y., & Jan, Y. N. (2008). Drosophila egg-laying site selection as a system to study simple decision-making processes. Science, 319(5870), 1679–1683. 10.1126/SCIENCE.1151842/SUPPL_FILE/YANG_SOM.PDF

Zhang, B., Segraves, K. A., Xue, H.-J., Nie, R.-E., Li, W.-Z., & Yang, X.-K. (2015). Adaptation to different host plant ages facilitates insect divergence without a host shift. Proceedings of the Royal Society B: Biological Sciences, 282(1815), 20151649. 10.1098/rspb.2015.1649

